# Discovery of a novel *Trichuris* species, *T. incognita*, with low sensitivity to albendazole-ivermectin treatment in Côte d’Ivoire patients, identified through fecal DNA metabarcoding

**DOI:** 10.1101/2024.07.04.602123

**Authors:** Abhinaya Venkatesan, Rebecca Chen, Max Baer, Pierre H. H. Schneeberger, Brenna Reimer, Eveline Hürlimann, Jean T Coulibaly, Said M Ali, Somphou Sayasone, John Soghigian, Jennifer Keiser, John Stuart Gilleard

## Abstract

Albendazole-ivermectin combination therapy is a promising alternative to benzimidazole monotherapy for *Trichuris trichiura* control. We used fecal DNA metabarcoding to genetically characterize *Trichuris* populations in patients from a clinical trial showing lower albendazole-ivermectin efficacy in Côte d’Ivoire (ERR below 70%) than in Lao PDR and Tanzania (ERR above 98%). ITS-1 and ITS-2 rDNA metabarcoding revealed the entire Côte d’Ivoire *Trichuris* population was phylogenetically distinct from *T. trichiura* in Lao PDR and Tanzania, and more closely related to the porcine parasite *Trichuris suis*. Complete mitogenomes of eight adult *Trichuris* from Côte d’Ivoire confirmed their species-level differentiation. Corresponding *Trichuris* ITS-1 and ITS-2 sequences in NCBI from individual human patients in Cameroon and Uganda and three captive non-human primates suggest this newly recognized species, *T. incognita*, is distributed beyond Côte d’Ivoire and has zoonotic potential. Further work is needed to assess the impact of this *Trichuris* species on soil transmitted helminth control programs.

## Introduction

*Trichuris trichiura* is a Soil Transmitted Helminth (STH) infecting 465 million people globally, primarily in middle and low-income countries (*1*). Infections are most prevalent in children, with moderate to severe infections resulting in chronic dysentery, diarrhea, and stunted growth (*2*). Control is largely dependent on Preventive Chemotherapy in high-risk endemic regions using annual or biannual albendazole and mebendazole administration (*3,4*). Benzimidazole efficacy against *T. trichiura* is generally low; a meta-analysis comparing 38 clinical trials reported egg reduction rates (ERR) and cure rates (CR) against *T. trichiura* of between ∼50% and ∼30% for albendazole, and ∼66% and ∼42% for mebendazole (*5*). However, albendazole-ivermectin combination therapy has demonstrated improved efficacy against *T. trichiura (6-11*). The WHO has consequently added this drug combination to the Essential Medicine List (EML) for soil transmitted helminths (*3, 11*).

A double-blind, parallel-group, phase 3 randomized, controlled clinical trial recently showed the expected high efficacy of albendazole-ivermectin combination therapy against *Trichuris* in Lao PDR, and Pemba, Tanzania, (ERR 99% and 98%, respectively) but an unexpectedly low efficacy in C□te d’Ivoire (ERR 70%) (*12*). Here, we have applied fecal DNA metabarcoding to investigate the genetics of the *Trichuris* populations from this clinical study.

Community-scale genetic analysis of STH is challenging because harvesting large numbers of parasites from patients, whether adult worms from expulsion studies or parasite eggs from feces, is labor intensive and can be logistically difficult. DNA metabarcoding applied directly to fecal DNA offers a powerful alternative approach. However, *Trichuris* eggs are robust and difficult to disrupt within fecal matter, which, together with fecal PCR inhibitors, presents a challenge for PCR and metabarcoding applied directly to fecal DNA (*13*). Multiple studies have aimed to improve the amplification of *Trichuris* DNA from human fecal samples by including a bead-beating step to mechanically disrupt the eggs (*14-17*).

In this study, we have used a DNA extraction protocol combining multiple freeze-thaw cycles and mechanical disruption to apply ITS-1 and ITS-2 rRNA fecal metabarcoding to genetically characterize *Trichuris* populations. Our results demonstrate that the albendazole-ivermectin refractory *Trichuris* population in the analyzed Côte d’Ivoire patients is comprised of a previously unrecognized species, *T. incognita*. A small number of ITS-2 sequences are present in NCBI GenBank that correspond to this species: two from human patients (from Cameroon and Uganda) and three from captive non-human primates, suggesting that this parasite is not unique to Côte d’Ivoire and may be zoonotic.

## Materials and Methods

### DNA extraction

#### Fecal samples

Fecal samples preserved in ethanol were collected from patients enrolled in a previously reported clinical trial (*12*). DNA was prepared from pre-treatment fecal samples from 22, 36, and 29 patients from C□te d’Ivoire, Lao PDR, and Pemba, respectively. *Trichuris* fecal egg counts ranged between 91-1151 epg, (Supplementary Table S1). Approximately 250 mg of feces in 90% ethanol were washed three times by 12,000*xg* centrifugation (1:2 ratio of feces: molecular-grade water) before three cycles of snap freezing in liquid nitrogen and 15 minutes of heating at 100°C with shaking at 750 rpm followed by vigorous bead beating for three minutes (Mini Bead Beater MBB-96, Thomas Scientific). DNA was then extracted using the QiAmp PowerFecal DNA extraction kit (Qiagen, Cat: 12830-50) (Appendix A1).

#### Adult worms

Whole worms were picked directly from patient stool samples following anthelmintic treatment during an expulsion study in Côte d’Ivoire (*18*). Worms were washed with sterile PBS and stored in absolute ethanol. DNA was extracted from separated worm heads using a DNeasy Blood & Tissue Kit (Qiagen, Cat: 69504).

### Fecal DNA metabarcoding

PCR primers were designed to amplify the following *Trichuris* genetic markers from fecal DNA: ITS-1 (733 bp), ITS-2 (592 bp), *cox-1* (430 bp), *nad-1* (470 bp), and *nad-4* (446 bp) (Supplementary Table S1). PCR reactions comprised 5 μL KAPA HiFi buffer (5X), 0.75 μL Illumina-adapted forward primer (10 μM), 0.75 μL Illumina-adapted reverse primer (10 μM), 0.75 μL dNTPs (10 μM), 0.5 μL KAPA HiFi Hotstart polymerase (0.5U), 0.1 μL bovine serum albumin (Thermo Fisher Scientific), 15.15 μL molecular grade water, and 2 μL of the extracted fecal DNA. Thermocycling conditions were 95□ for 3 min, followed by 40 cycles of 98□ for 20s, 65□ (ITS-1), 64□ (ITS-2, *nad-1, nad-4*), 60□ (*cox-1*), and 63□ (β-tubulin) for 15 s, 72□ for 30 s, followed by 72□ for 2 min. Library preparation from amplicons was performed as previously described (*19*). Amplicons were sequenced on an Illumina MiSeq using the V3 (2 × 300 bp) sequencing chemistry for ITS-2, *nad-1*, and *nad-4*, and V2 (2 × 250 bp) sequencing chemistry for ITS-1 *and cox-1*.

Quality filtering of paired-end reads and primer removal were performed using Cutadapt v3.2 (*20*), and the adapter-trimmed paired-end reads were analyzed using DADA2 as previously described (*21, 22*). Forward reads were trimmed to 240 bp for ITS-1, ITS-2, *cox-1, nad-1, nad-4* amplicons, whereas the reverse reads were trimmed to 190 bp, 220 bp, 230 bp, 220 bp, and 220 bp for the ITS-1, ITS-2, *cox-1, nad-1*, and *nad-4*, amplicons, respectively. Reads shorter than 50 bp or with an expected error rate of greater than 1, 3, 1, 2, and 2, nucleotides for forward reads and greater than 2, 5, 2, 5, and 5 for reverse reads of ITS-1, ITS-2, *cox-1, nad-1, and nad-4* amplicons, respectively, were removed. Amplicon Sequence Variants (ASVs) were aligned to reference sequences (Supplementary Table S2) using the MAFFT tool for multiple sequence alignment (*23*) and off-target ASVs were removed. Manual correction of sequence alignments was performed using Geneious v10.0.9. Only samples with a total read depth >1000 mapped reads and ASVs present at >0.1 % in the population overall and at least 200 reads in an individual sample were included in the final analysis.

### Phylogenetic and population genetic analysis of metabarcoding data

Multiple sequence alignment of *Trichuris* ITS-1, ITS-2, *cox-1, nad-1*, and *nad-4* reference sequences from GenBank (Supplementary Table S2) and the ASVs from amplicon sequencing was performed using the MAFFT tool for multiple sequence alignment. Following manual correction for indels in Geneious v10.0.9, phylogenetic analysis was performed using the Maximum Likelihood (ML) method.

Statistical parsimony haplotype networks were constructed from the corrected alignments of the ASVs using the pegas() package in R (*24*). These networks were then visualized and annotated on Cytoscape v3.9.1 (*25*).

Following ASV filtering and alignment, haplotype distribution bar charts were generated in R using the ggplot2() package (*26*). The Shannon-Wiener index for alpha diversity was calculated using the vegan() package in R (*27*) and plotted for each population using ggplot2(). Pairwise t-test was used to determine significance of differences in the alpha diversity between populations. For beta diversity analysis, a Bray-Curtis dissimilarity matrix and Jaccard distance were used to calculate differences in relative abundance and presence or absence of ASVs among the samples, respectively. Principal Coordinate Analysis (PCoA) was performed using the vegan() package in R, and the plots were generated using ggplot2(). Nucleotide diversity and ASV heterozygosity were calculated using the pegas() package in R, whereas pairwise FST calculations and significance testing were performed in Arlequin v3.5 (*28*) using 1000 permutations.

### Adult worm whole genome sequencing and analysis

Adult worm Illumina sequencing libraries were prepared using the NEBNext® Ultra™ II FS DNA Library Prep Kit (New England Biolabs, Cat: E7805L) and DNA concentrations measured with the dsDNA High Sensitivity Assay kit on a Qubit 4 Fluorometer (both Invitrogen, USA) using 2µL of DNA extract as input. Fragment size was checked via 1% gel electrophoresis and libraries were sequenced using 2×150 bp paired-end chemistry on an Illumina MiSeq.

Following whole genome sequencing, *fastp (29*) was used to filter reads below Phred quality score of 15 and trim adapters from paired end reads. Next, GetOrganelle (*30*) was used to assemble mitochondrial genomes and extract ITS-1 and ITS-2 sequences from each sample. For assembling mitochondrial genomes, GetOrganelle was run with flags “-R 10”, “-k 21, 45, 65, 85, 105” and a default animal mitochondrial genome database seed (-F animal_mt). Mitochondrial genomes were then annotated in Geneious v10.0.9 using the “Annotate From” function with *Trichuris* reference genomes from GenBank (Supplementary Table S2), with a threshold of 90% similarity. For extracting ITS-1 and ITS-2 sequences, two *Trichuris* sequences from GenBank were used as seeds (accessions AM993005 and AM993010), -F was set to “anon”, and a target size of 1350 was used.

Additional mitochondrial genomes representing 13 described species of *Trichuris*, several undescribed lineages of *Trichuris*, and *Trichinella* outgroups were downloaded from GenBank or retrieved from Doyle et al., 2022 (*13*) (Supplementary Table S2). Protein coding sequences for the genes *ATP6, cox-1, cox-2, cox-3, cytb, nad-1, nad-2, nad-3, nad-4, nad-4L, nad-5*, and *nad-6* were extracted from all 45 mitogenomes and aligned separately with MAFFT. Pairwise nucleotide identity was calculated in Geneious v10.0.9. IQ-TREE 2 (*32*) was used to estimate the maximum likelihood phylogeny from the concatenated alignment of the 12 protein-coding genes, with a separate substitution model estimated per gene (flag -m MFP). Support for branches in the maximum likelihood topology was assessed with both Ultrafast Bootstrap (1000 replicates; -B 1000) and an SH-like aLRT branch test (--alrt 1000). The resulting maximum likelihood phylogeny was visualized in FigTree (*33*) and rooted on the branch leading to *Trichinella*.

## Results

### ITS1 and ITS2 rDNA metabarcoding suggests the *Trichuris* population in Côte d’Ivoire samples is entirely comprised of a species distinct from *T. trichiura* present in Pemba and Lao PDR

The 733 bp ribosomal ITS-1 region was amplified and deep sequenced from 21, 34, and 29 fecal samples, and the 592 bp ITS-2 region from 15, 26, and 23 fecal samples from patients in C□te d’Ivoire, Lao PDR, and Pemba Island, respectively. ITS-1 and ITS-2 paired-end reads were merged to produce 461bp and 457bp sequences with average mapped read depths of 17,246 (range: 2,205 - 28,402) and 7,660 (range: 1,172 - 18,027), respectively.

While many ASVs were shared between the *Trichuris* populations in Lao PDR and Pemba, there were no ASVs shared with the C□te d’Ivoire populations (Figure 1A). Pairwise FST values were much higher between C□te d’Ivoire and Lao PDR, and C□te d’Ivoire and Pemba than between Lao PDR and Pemba (Table 1). The ITS1 and ITS-2 ASVs present in the C□te d’Ivoire *Trichuris* population were genetically very divergent from those found in the Lao PDR and Pemba populations, being separated by 145/461 and 196/457 nucleotide differences for ITS-1 and ITS-2, respectively (Figure 1B).

**Table 1:** Pairwise FST comparison among different *Trichuris* populations.

| ITS-1 |  |  | ITS-2 |  |  |
| --- | --- | --- | --- | --- | --- |
| Region | Lao PDR | Pemba | Region | Lao PDR | Pemba |
| Côte d'Ivoire | 0.248 | 0.273 | Côte d'Ivoire | 0.300 | 0.336 |
| Lao PDR |  | 0.017 | Lao PDR |  | 0.015 |

**Figure 1:**
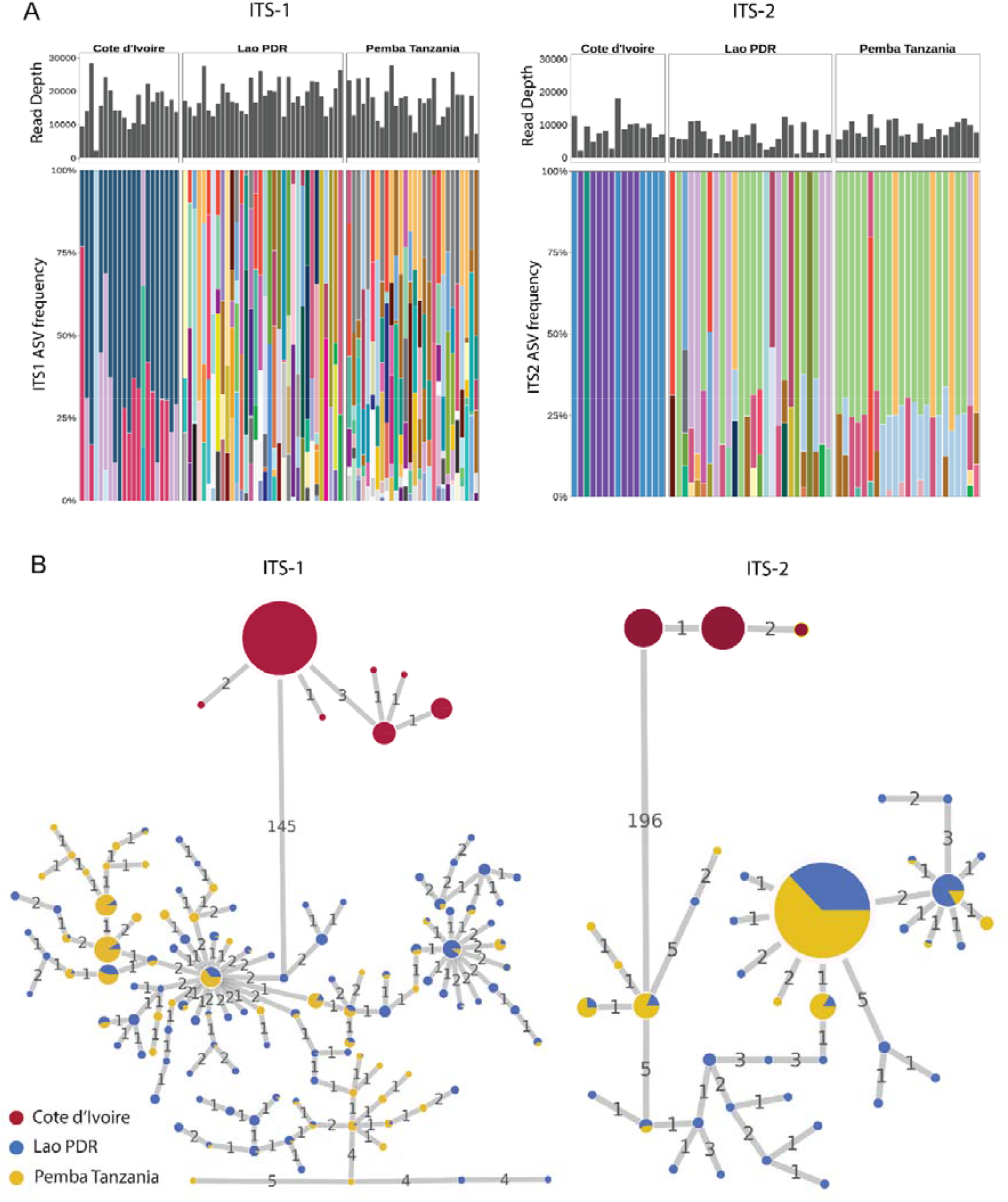
Panel A: Trichuris ITS-1 and ITS2 rDNA ASVs generated by amplicon sequencing from fecal samples of Côte d’Ivoire, Lao PDR, and Pemba patients. Panels A and B showing the relative frequencies of ASVs generated by amplicon sequencing of the Trichuris ITS-1 and ITS-2 loci, respectively, from samples in Côte d’Ivoire, Lao PDR, and Pemba. The histogram at the top of each panel shows the mapped read depth for each sample, and the bar plots below represent the relative frequencies of the ASVs present in each sample from the three regions. **Panel B: Haplotype networks of the ITS-1 and ITS2 rDNA ASVs generated by amplicon sequencing from fecal samples of Côte d’Ivoire, Lao PDR, and Pemba patients**. The figure shows statistical parsimony haplotype networks of ASVs for the Trichuris ITS-1 (A) and ITS-2 (B) loci generated by amplicon sequencing from fecal samples of Côte d’Ivoire, Lao PDR, and Pemba patients. Each circle represents an ASV, the size of which represents its frequency. The numbers on the connecting lines indicate the number of nucleotide differences between adjacent haplotypes. ASVs are colored based on which region they are located.

Phylogenetic analysis of the *Trichuris* ITS-1 and ITS-2 ASVs from the three regions had broadly congruent results for the two markers (Figure 2). All *Trichuris* ASVs from C□te d’Ivoire form a separate phylogenetic clade to those from Lao PDR and Pemba, clustering with *Trichuris* reference sequences from human patients in Cameroon and Uganda, and from captive non-human primates from Italy, Czech Republic, and Uganda. This clade is more closely related to the clade containing *Trichuris suis* than that containing ASVs from Lao PDR and Pemba for both markers (Figure 2). Côte d’Ivoire *Trichuris* ITS-2 and ITS-1 ASVs shared 65% and 80% identity with the *T. suis* sequences but only 44% and 56% identity with Lao PDR and Pemba *Trichuris* ASVs, respectively.

**Figure 2:**
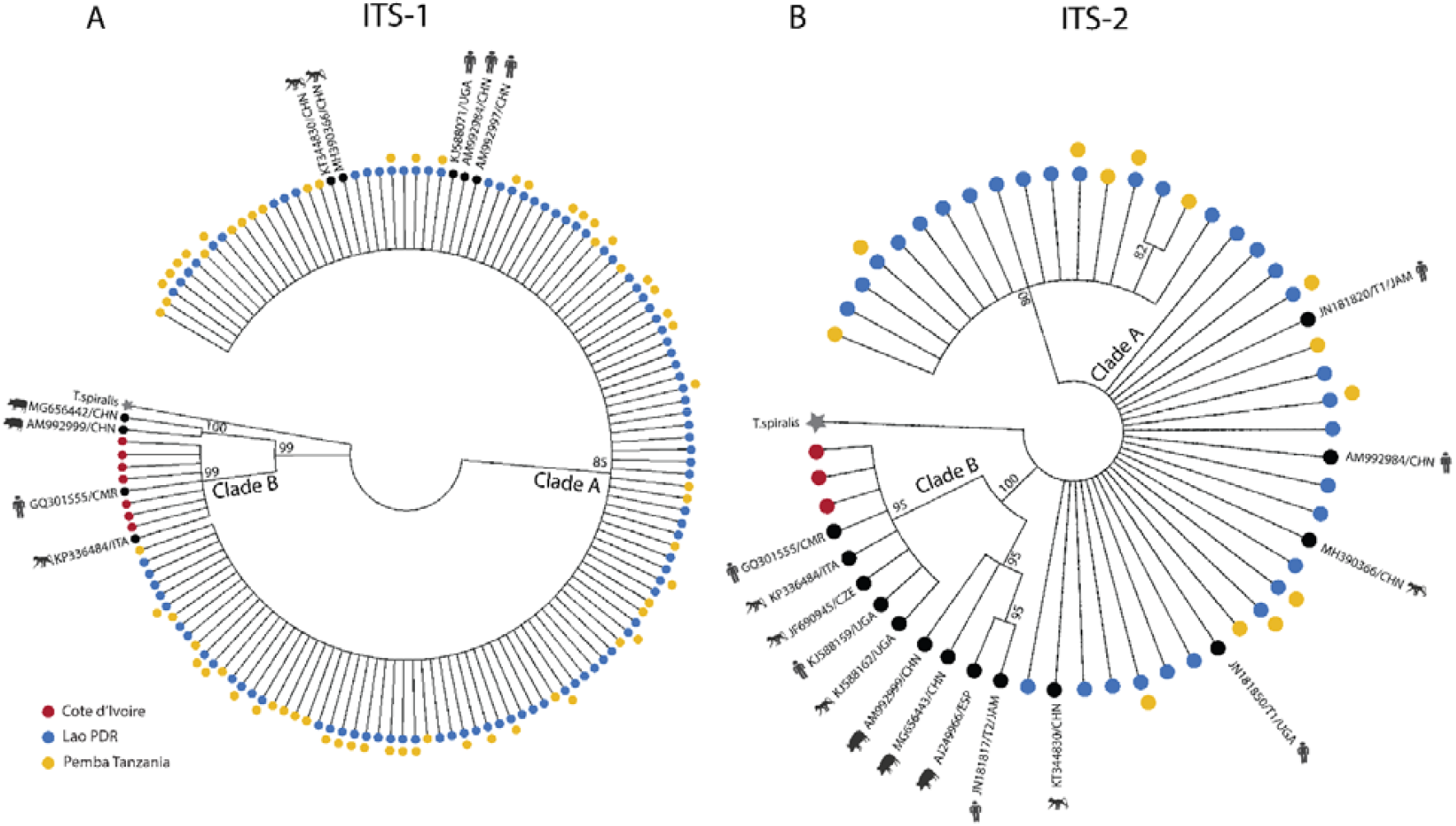
Maximum likelihood phylogenetic tree of the ITS-1 and ITS2 rDNA ASVs generated by amplicon sequencing from fecal samples of Côte d’Ivoire, Lao PDR, and Pemba patients. Maximum likelihood trees of the *Trichuris* ITS-1 (A) and ITS-2 (B) ASVs from Côte d’Ivoire, Lao PDR, and Pemba, as well as additional *Trichuris* reference sequences available from pigs, humans, and non-human primates on GenBank. *Trichinella spiralis* (GenBank accession: KC006432) is used as the outgroup. Each tip of the tree is an ASV or a sequence from GenBank and the color represents which regions the ASV is present in. For the ITS-1 and ITS-2 sequences, the Kimura 80 (K80) and general time reversible (GTR) models were chosen as the best nucleotide substitution models, respectively, whereas the gamma-distributed rate variation (GTR+G) general time reversible model was chosen for the mitochondrial cox-1, nad-1, and nad-4 markers. These models were chosen using jModeltest v2.1.10 (Darriba D et al., 2012). ML trees were constructed using PhyML v3.3 (Guindon S et al., 2010) with 100 bootstrap replicates using *Trichinella spiralis* as the outgroup (GenBank accession: KC006432 for the ITS-1 and ITS-2 sequences, NC002681 for the mitochondrial cox-1, nad-1, and nad-4 sequences). Using MEGA v11 (Tamura K et al., 2021), the ML trees were condensed to only display branches with consensus support >80%.

ASV heterozygosity and nucleotide diversity were much higher in Pemba and Laos, with much higher alpha diversity than in C□te d’Ivoire for both ITS-1 and ITS-2 (Figure 3, Supplementary Table S4).

**Figure 3:**
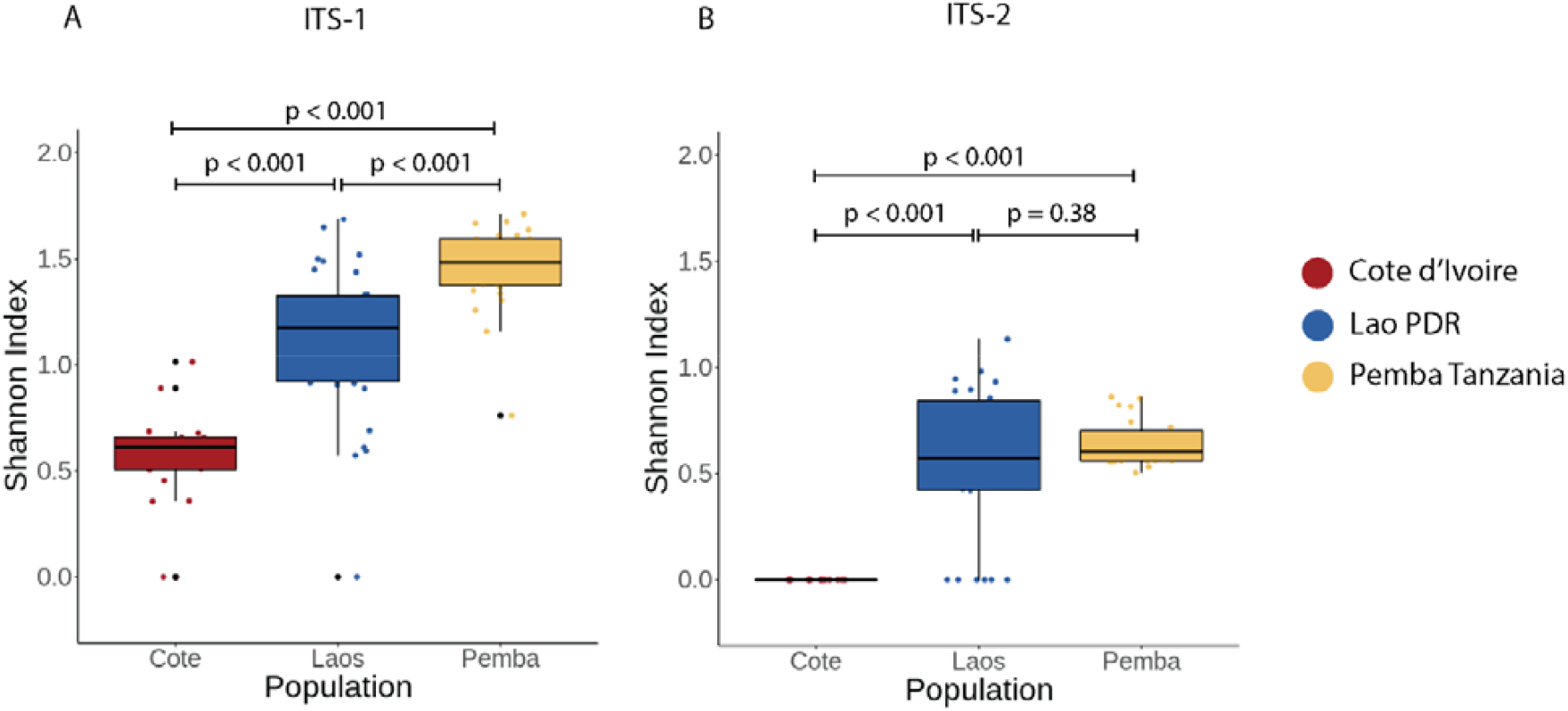
Alpha diversity of ITS-1 and ITS2 rDNA ASVs generated by amplicon sequencing from fecal samples of Cote d’Ivoire, Lao PDR, and Pemba patients. Panels A and B show the Shannon-Wiener Index, representing alpha diversity, for the ASVs generated by amplicon sequencing of the *Trichuris* ITS-1 and ITS-2 loci, respectively, from patient fecal samples in Côte d’Ivoire, Lao PDR, and Pemba. The p-values from the pairwise t-tests are indicated between the regions.

### Mitochondrial *cox-1, nad-1*, and *nad-4* metabarcoding confirms Lao PDR and Pemba

#### *Trichuris* populations are *T. trichiura* and reveals population sub-structuring

Species-specific primers designed against *T. trichiura* reference sequences (GenBank accessions: NC_017750, GU385218, AP017704, and KT449825) successfully generated mitochondrial *cox-1, nad-1*, and *nad-4* amplicons for metabarcoding from the majority of Lao PDR and Pemba fecal samples but not from any C□te d’Ivoire samples, suggesting primer site sequence polymorphism of this latter *Trichuris* population (Figure S1).

Mitochondrial *cox-1, nad-1*, and *nad-4* ASVs generated from Lao PDR and Pemba revealed high alpha diversity (Figures S2 and S3), and all ASVs clustered phylogenetically with *T. trichiura* reference sequences, supporting their species identity (Figure S4). Although pairwise FSTs between the Pemba and Lao PDR *T. trichiura* populations were low (0.024, 0.002, and 0.142 for *cox-1, nad-1*, and *nad-4*, respectively), multi-dimensional metric beta diversities and haplotype networks indicated some geographical sub-structuring between the regions, particularly for the *nad-1*marker (Figures S5 and S6).

#### Analysis of complete mitogenomes from Côte d’Ivoire adult worms support their separate species status

Since the primers designed against *T. trichiur*a mitochondrial reference sequences did not yield amplicons from the C□te d’Ivoire fecal DNA samples, we extracted and assembled complete mitochondrial genomes from whole genome sequencing data (WGS) obtained from eight adult *Trichuris* worms as part of a separate anthelmintic expulsion study in the C□te d’Ivoire study population (*18*). Average mitochondrial genome coverage was 2,541X and the eight assembled mitochondrial genomes ranged in size from 14,253bp to 14,663bp (average: 14,338.13 bp), slightly larger than *T. trichiura* reference genomes retrieved from GenBank (range: 14,046-14,091, n=6) but similar to those of *T. suis* (range: 14,436-14,521, n=3). Phylogenetic reconstruction of protein coding genes from *Trichuris* mitochondrial genomes indicated that the eight C□te d’Ivoire mitogenomes formed a single, well-supported clade, sister to *Trichuris* from Colobus monkeys (*31*) and more genetically distant from *T. trichiura* (Figure 4). Despite this phylogenetic similarity, adult worms sampled from C□te d’Ivoire were genetically distinct from Colobus monkeys and *T. suis*, with pairwise nucleotide identities from COX1 of 80.2% and 78.9% respectively, and 73.1% and 71.7% across all protein coding genes (Supplementary Table S5).

**Figure 4:**
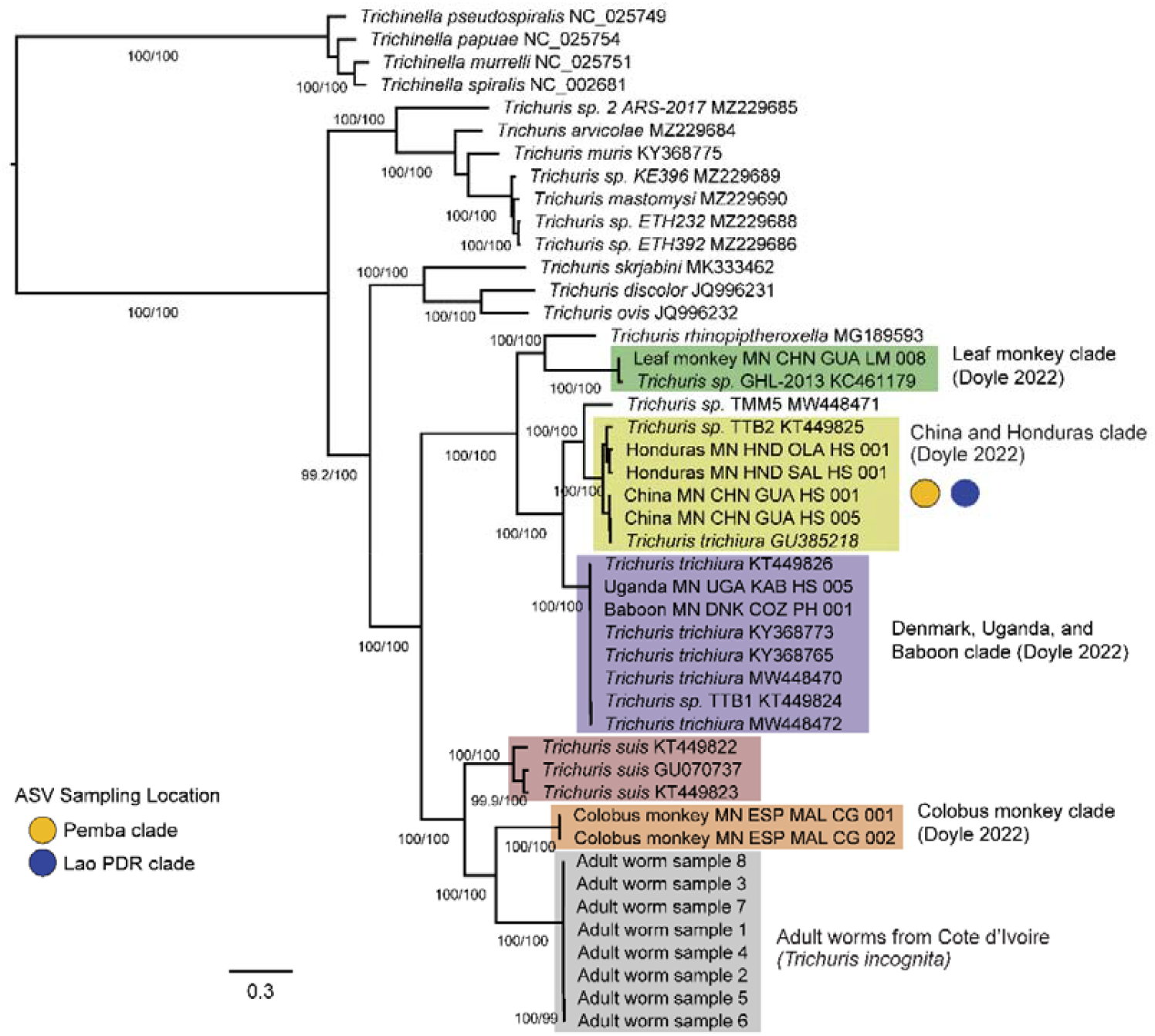
The maximum likelihood phylogeny of *Trichuris* species from twelve mitochondrial protein coding genes. The phylogeny was constructed in IQ-TREE with alignments from twelve protein coding genes from 45 mitochondrial genomes, including eight *T. incognita* samples obtained from expulsion studies of Cote d’Ivoire patients (*18*). Ultrafast bootstrap values and SH-like branch support values above 95/95 are shown as numbers on branches. Scale is in substitutions per site. Colored circles refer to amplicon sequencing variant groups and their sampling location, and are placed next to clades to indicate relatedness between these sequences and samples from the highlighted clades, based on other analyses.

The ITS-1 and ITS-2 rDNA sequences recovered from the WGS of the eight adult worms were aligned to the ASVs generated from the fecal metabarcoding. Sequence identities ranged from 93.5% to 99.8%, indicating that the same species of *Trichuris* was sampled in the adult worm whole genome sequencing as the fecal stool metabarcoding (Figures S7 and S8)

## Discussion

Co-administration of benzimidazoles with macrocyclic lactones provides much higher efficacy against *T. trichiura* than monotherapy with either drug class (*5-10*). A double-blind, parallel-group, phase 3, randomized, controlled trial was recently conducted to compare the efficacy of albendazole-ivermectin combination with albendazole monotherapy in C□te d’Ivoire, Lao PDR, and Tanzania. While the combination therapy had much higher efficacy than albendazole monotherapy in Lao PDR and Tanzania, the efficacy in C□te d’Ivoire was much lower and comparable to monotherapy (*12*). Here we have used short-read metabarcoding of several taxonomic markers to compare the genetics of the *Trichuris* populations in fecal samples from multiple patients at each of the three study sites. Analysis of the ITS-1 and ITS-2 ASVs revealed a genetically divergent *Trichuris* population in C□te d’Ivoire that did not share any ASVs with the samples in Lao PDR or Pemba (Figure 1). Pairwise FST calculations and haplotype networks revealed significant population differentiation in the C□te d’Ivoire *Trichuris* population.

The ASVs of both ITS-1 and ITS-2 from our study clustered into two distinct phylogenetic clades: Clade A and Clade B (Figure 2). All the ASVs from the Lao PDR and Pemba patients belong to Clade A which corresponds to the majority of previously described reference sequences for *T. trichiura* in public databases (*34-41*). A varied previous nomenclature has used for these *Trichuris* sequences within our Clade A from in the public databases and literature; Subclade DG (*35*), Group 1 (*36*), Clade DG (*39*), Clade 2 (*40*), or Subgroup 1 (*41*). In contrast, all the *Trichuris* ASVs generated from Côte d’Ivoire patients fall into a second monophyletic clade (Clade A) much closer to reference sequences from the porcine parasite *T. suis* than to human *T. trichiura* (Figure 2). Nevertheless, Clade B is distinct from *T. suis* and is more closely related to a small number of reference sequences from humans and non-human primates previously deposited in public databases (*35-36, 39-41*). Specifically, one human patient from Cameroon, one human patient from Uganda (*39*) and several additional reference sequences from red colobus monkey in Uganda, captive vervet monkey in Italy, and captive lion-tailed macaque in Czech Republic (*35-36*) (Figure 2). The nomenclature previously used in the literature for this sequence group is also varied; Subclade CA (*35*), Group 3 (*36*), CP-GOB (*39*), Clade 1 (*40*), or Subgroup 5 (*41*).

Primers designed against *T. trichiura* mitochondrial *cox-1, nad-1* and *nad-4* reference sequences consistently generated PCR amplicons from the samples in Lao PDR and Pemba, but not from Côte d’Ivoire. The PCR amplification failure from the Côte d’Ivoire samples is unsurprising given the genetic divergence of this *Trichuris* population from *T. trichiura*. Although, some population genetic sub-structuring was apparent between the Lao PDR and Pemba *Trichuris* populations from the mitochondrial DNA metabarcoding, all ASVs from these two sites clustered with human-derived *T. trichiura* reference sequences from the public databases, including sequences from China, Japan, and Honduras, and with *Trichuris* sequences from captive baboons in the USA (*31*) (Figure S4). This, together with the ITS1 and ITS2 ASV data, shows the albendazole-ivermectin sensitive *Trichuris* populations from Lao PDR and Pemba are entirely comprised of *T. trichiura*.

To investigate the mitochondrial phylogeny of the Côte d’Ivoire *Trichuris* sequences, we assembled the complete mitochondrial genomes from eight adult *Trichuris* worms collected from an expulsion study in Côte d’Ivoire (*18*). Phylogenetic analysis of the eight assembled mitochondrial genomes, in conjunction with previously released 37 mitogenomes produced a similar outcome to the ITS marker analysis. These mitochondrial genomes are more closely related to *Trichuris* from Colobus monkeys (*31*) and *T. suis* than they are to *T. trichiura* in the public databases (Figure 4). Despite the phylogenetic relationship of the Côte d’Ivoire *Trichuris* to these *Trichuris* from other animals, the genetic distance between these lineages, as represented in our phylogenetic analysis and in pairwise nucleotide identity analysis (Supplementary Table S5), indicate that we have sampled a new species of *Trichuris*. The reference genome of this previously unrecognized *Trichuris* species, proposed *Trichuris incognita*, has been characterized concurrent to this study (*18*). We summarize the overall phylogenetic relationships of *T. trichiura, T. suis*, and *T. incognita* in Figure 5.

**Figure 5:**
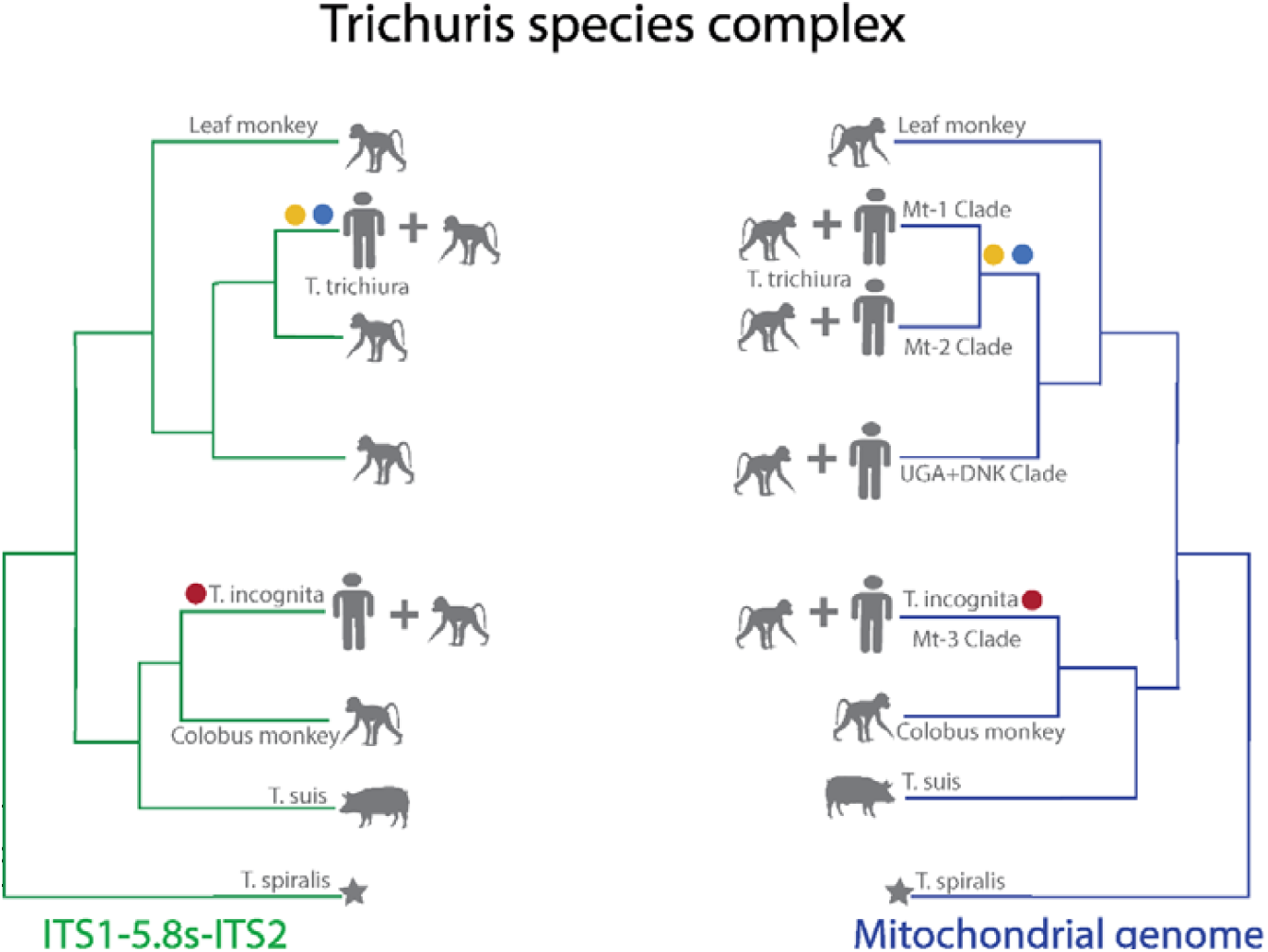
Schematic representation of the *Trichuris* species complex. Schematic representation of the phylogenetic relationships of *Trichuris* infecting humans and non-human primates adapted from previously published studies and current findings on the analysis of ribosomal and mitochondrial markers. The figure shows two major clades of *Trichuris* in the ribosomal DNA phylogeny and mitochondrial DNA phylogeny infecting

It is noteworthy that the site in Côte d’Ivoire has the longest record of consistent use of albendazole-ivermectin treatment in MDA campaigns aimed at combating lymphatic filariasis. (more than two decades (*12, 42*). This contrasts with the situation in Pemba, where lymphatic filariasis MDA was stopped many years ago, or in Laos, where ivermectin has not been previously used (*12*). It is possible that this treatment history has provided a selective advantage for *T. incognita* in Côte d’Ivoire due to an inherently lower sensitivity to this drug combination compared with *T. trichiura*.

In spite of the studied Côte d’Ivoire samples having higher fecal egg counts (mean epg 350, range 110 - 1151), than samples in Lao PDR (mean epg 147, range 55 - 700) and Pemba (mean epg 202, range: 51 – 803), the Côte d’Ivoire *T. incognita* population had extremely low ITS-1 and ITS-2 alpha diversity relative to the *T. trichiura* populations (Figure 3, Supplementary Table S4). It is unlikely that this is explained by higher allelic dropout for *T. incognita* than *T. trichiura*, since the primers used have 100% identity with the ITS-1 and ITS-2 sequences of *T. suis* reference sequences). Low genetic diversity of the *T. incognita* population may indicate that this species has relatively recently adapted to human hosts, and its presence in non-human primates may be consistent with this hypothesis (*43, 44*). Alternatively, it could be the result of selective pressure due to the long history of albendazole-ivermectin MDA in the region.

In conclusion, we have identified a previously unrecognized human *Trichuris* species, *T. incognita*, that is more closely related to *T. suis* than to *T. trichiura* and is less responsive to albendazole-ivermectin combination therapy. This work demonstrates how fecal DNA metabarcoding can be used to characterize human helminth populations at the community level. The application of fecal DNA metabarcoding using ITS as well as other markers will be a powerful method to investigate the geographical distribution, treatment efficacy, and control strategies for this newly recognized *Trichuris* species.

