## Supplementary Information for "Discovery of a novel *Trichuris* species, *T. incognita*, with low sensitivity to albendazole-ivermectin treatment in Côte d’Ivoire patients, identified through fecal DNA metabarcoding" (from another version of this preprint: version 2, 2024-07-07)

### Supplementary Figures

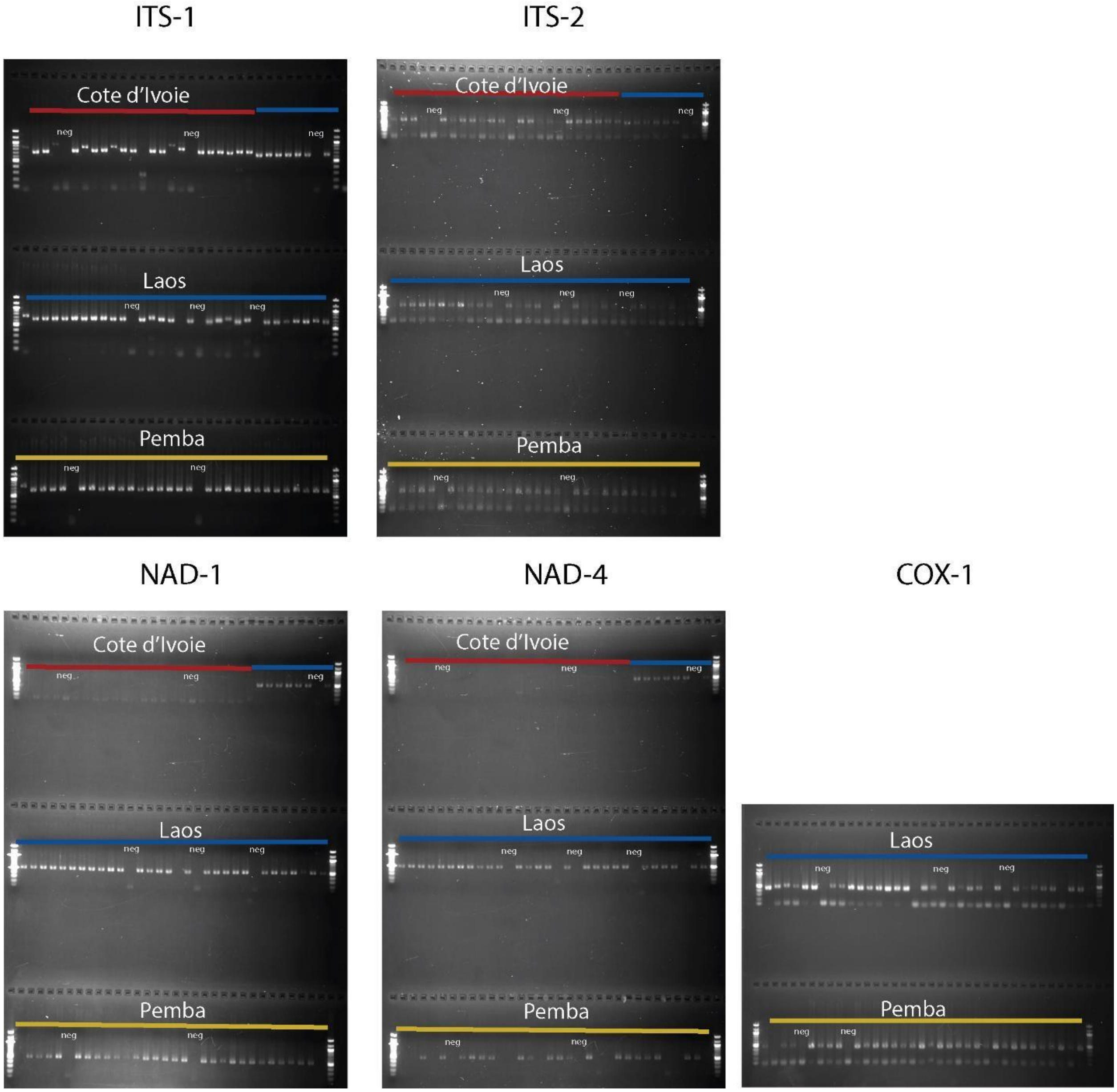


**Figure S1: Gel electrophoresis results from PCR amplicon generation**

Image of the gel electrophoresis results of the PCR amplification outcomes for the ITS-1, ITS-2 and mitochondrial *cox-1, nad-1,* and *nad-4* markers from patient fecal samples from Côte d'Ivoire, Lao PDR, and Pemba. Species-specific primers designed against *T. trichiura* reference sequences (Genbank accession numbers: NC_017750, GU385218, AP017704, and KT449825) successfully generated mitochondrial *cox-1, nad-1*, and *nad-4* marker amplicons from 32, 35, and 34 of the 36 Lao PDR samples and 29, 28, and 19 from the 29 Pemba samples, respectively. No amplicons were generated from any of the 22 Côte d'Ivoire samples.

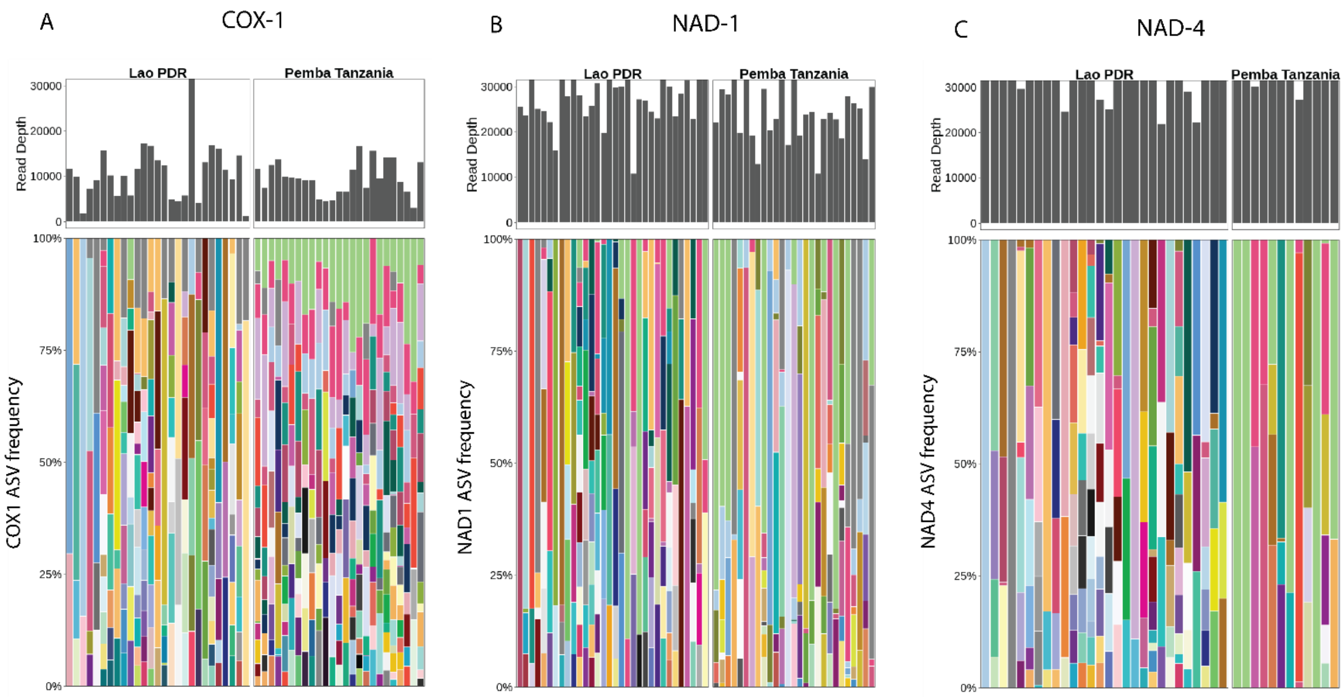


**Figure S2: ASVs generated by amplicon sequencing of *cox-1, nad-1,* and *nad-4* markers from fecal samples of Côte d'Ivoire, Lao PDR, and Pemba patients**

Panels A, B, and C are bar charts showing the relative frequencies of different ASVs of the *Trichuris* mitochondrial *cox-1, nad-1,* and *nad-4* markers, respectively, amplified from fecal samples from Lao PDR and Pemba. The histogram at the top of each panel shows the mapped read depth for each sample, and the bar plots below represent the relative abundance of the ASVs present in each sample from the three regions.

For Laos and Pemba, respectively, metabarcoding data was generated from 27 and 25 samples for the 430 bp *cox-1* marker, from 32 and 27 samples for the 470 bp *nad-1* marker and from 28 and 12 samples for the 446 bp *nad-4*. Paired-end reads from the three markers were merged and concatenated to produce average mapped read depths of 10,300 reads (range: 1,355 - 32,600), 26,490 reads (range: 10,791 - 56,330), and 36,350 reads (range: 21,936 - 55,346) for the *cox-1, nad-1,* and *nad-4* markers, respectively.

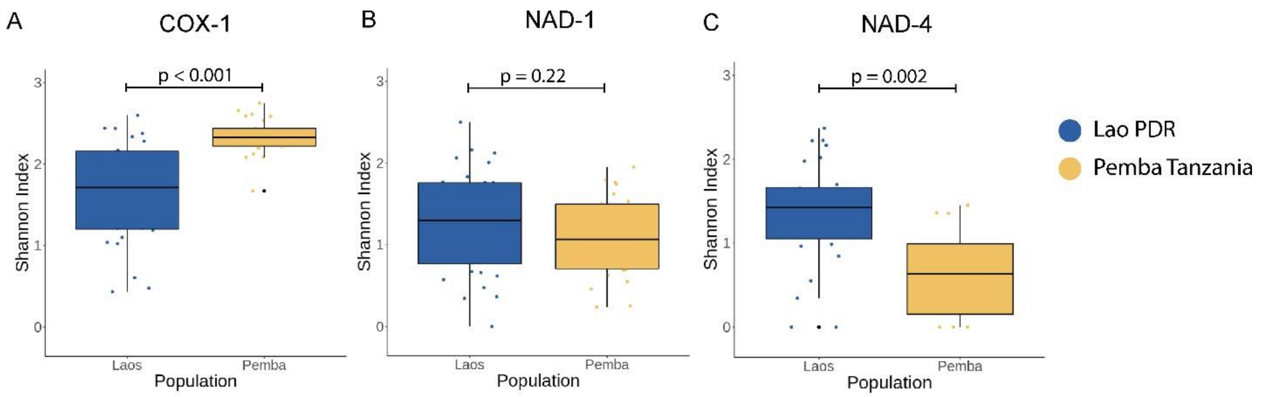


D

| Region | cox-1 | | | | nad-1 | | | | nad-4 | | | |
|---|---|---|---|---|---|---|---|---|---|---|---|---|
| | N | $N_{ASV}$ | $H_d$ | π | N | $N_{ASV}$ | $H_d$ | π | N | $N_{ASV}$ | $H_d$ | π |
| Laos | 33 | 166 | 0.98 | 0.038 | 31 | 121 | 0.98 | 0.012 | 28 | 147 | 0.98 | 0.014 |
| Pemba island | 29 | 129 | 0.96 | 0.022 | 27 | 106 | 0.93 | 0.034 | 13 | 27 | 0.80 | 0.008 |

**Genetic diversity in the mitochondrial DNA loci**: N = Number of samples sequenced; $N_{ASV}$ = Number of ASVs; $H_d$ = ASV heterozygosity; π = Nucleotide diversity

**Figure S3: Alpha diversity at the mitochondrial DNA locus**
Figures A, B, and C are box plots indicating the Shannon-Wiener Index to represent alpha diversity at the mitochondrial *cox-1, nad-1,* and *nad-4* loci, respectively, from samples in Lao PDR and Pemba. The p-values from the pairwise t-tests are indicated between the regions. The genetic diversity observed at the three mitochondrial loci are tabulated in D.

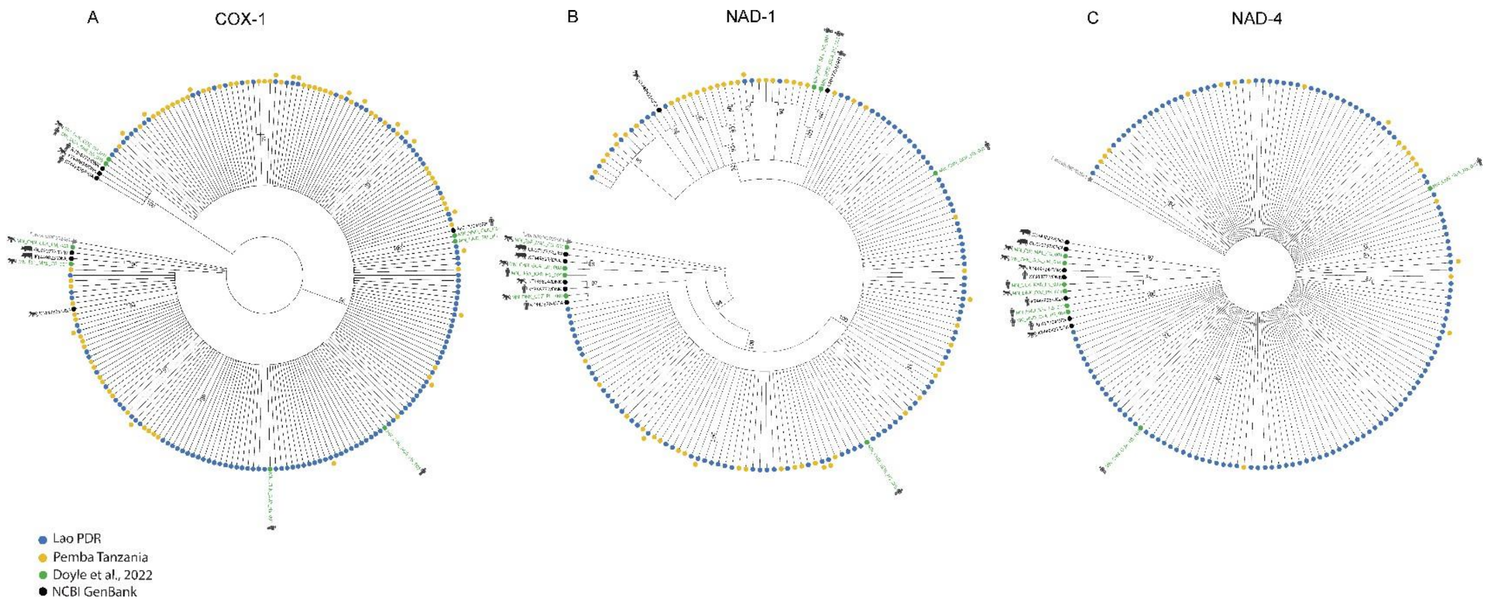


**Figure S4 : Maximum Likelihood phylogenetic tree of ASVs the mitochondrial *cox-1, nad-1,* and *nad-4* markers generated by amplicon sequencing from Lao PDR and Pemba patient fecal samples** Maximum likelihood trees of the mitochondrial *cox-1* (A), *nad-1* (B), and *nad-4* (C) marker ASVs amplified from fecal samples from patients in Lao PDR and Pemba as well as additional reference *Trichuris* sequences available from pigs, humans, and non-human primates on GenBank. *Trichinella spiralis* (GenBank accession: NC002681) is used as the outgroup. Each tip of the tree is an ASV or a sequence from GenBank and the color represents which regions the ASV is present in.

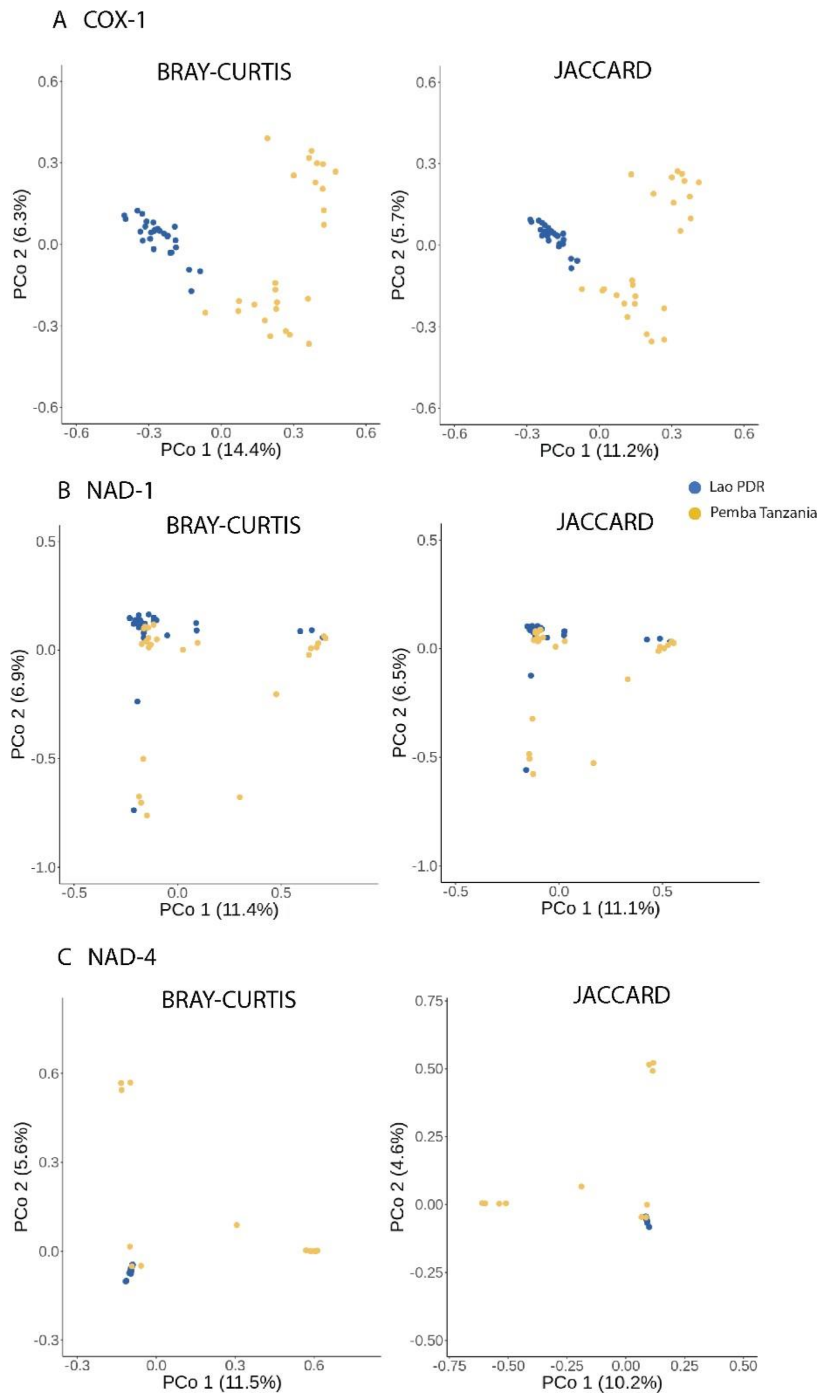


**Figure S5: Multi-dimensional metric analysis showing Beta diversity based on the *cox-1, nad-1,* and *nad-4* ASV data generated by amplicon sequencing from patient samples Lao PDR and Pemba the from at the mitochondrial DNA locus**

Figures A, B, and C are PCoA plots of the mitochondrial *cox-1, nad-1,* and *nad-4* loci, respectively, from samples in Lao PDR and Pemba. Multi-dimensional metric analysis was performed using two different methods: the Bray-Curtis dissimilarity matrix which calculates distances based on the abundance of ASVs, and the Jaccard index which calculates distances based on the presence or absence of ASVs. Each point is an individual patient sample, and the points are colored based on the region in which they are present.

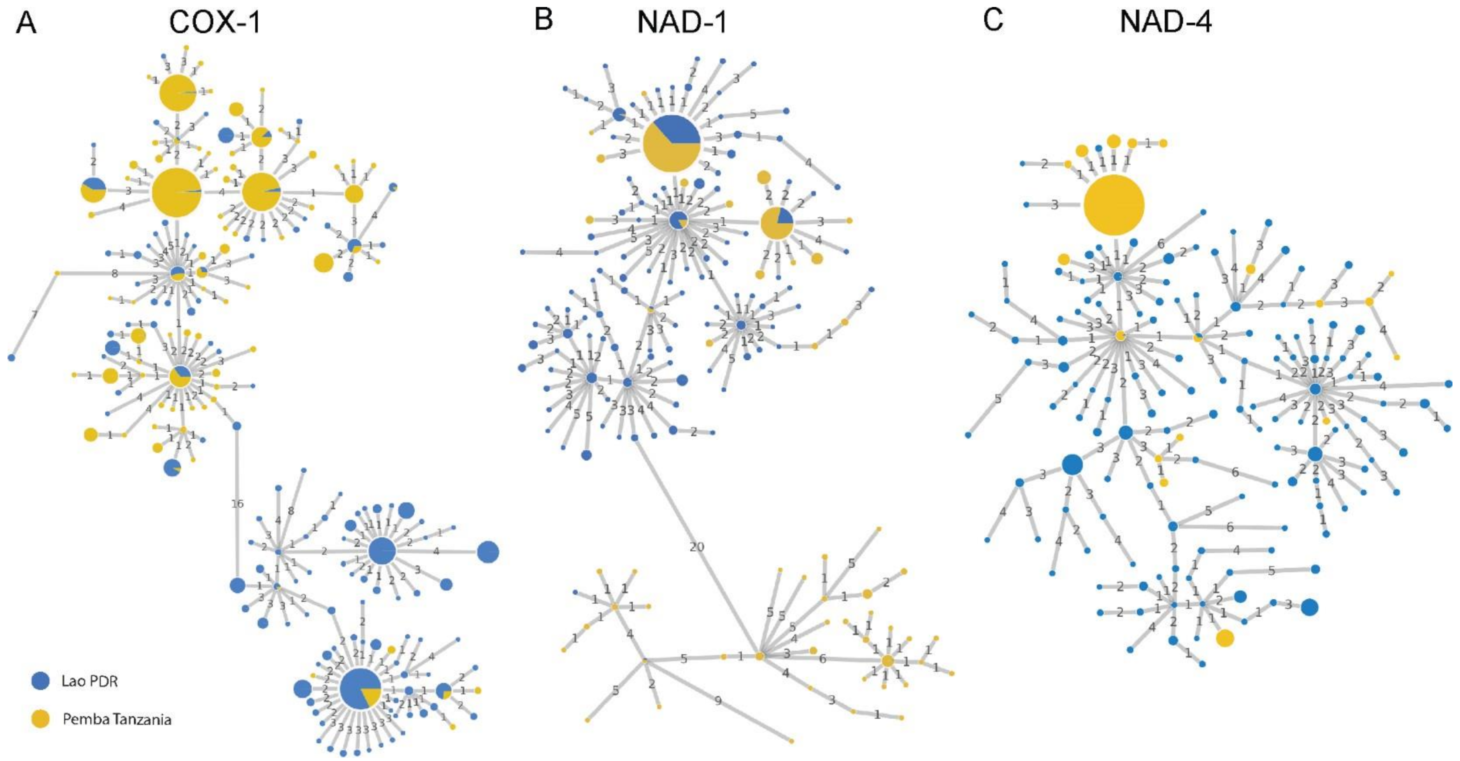


**Figure S6: Haplotype network of ASVs of the mitochondrial *cox-1, nad-1,* and *nad-4* markers generated by amplicon sequencing from Lao PDR and Pemba patient fecal samples**

The figure comprises statistical parsimony haplotype networks of ASVs for the *Trichuris* mitochondrial *cox-1* (A), *nad-1* (B), and *nad-4* (C) markers generated by amplicon sequencing from fecal samples from patients in Lao PDR and Pemba. Each circle represents an ASV, the size of which represents its frequency. The numbers on the connecting lines indicate the number of nucleotide differences between adjacent haplotypes. ASVs are colored based on which region they are present in. The haplotype networks show the major (high-frequency) ASVs differed between the two regions for all three markers and most of the rarer (low-frequency) ASVs were unique to either region.

A

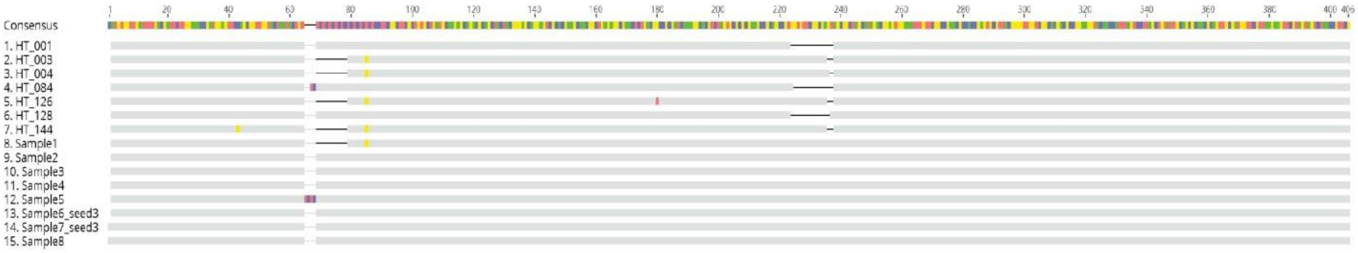


B

| | HT_001 | HT_003 | HT_004 | HT_084 | HT_126 | HT_128 | HT_144 | Sample1 | Sample2 | Sample3 | Sample4 | Sample5 | Sample6 | Sample7 | Sample8 |
|---|---|---|---|---|---|---|---|---|---|---|---|---|---|---|---|
| HT_001 | | 94.2 | 94.0 | 99.2 | 94.0 | 99.7 | 94.0 | 93.8 | 96.5 | 96.5 | 96.5 | 95.6 | 96.5 | 96.5 | 96.5 |
| HT_003 | 94.2 | | 99.7 | 94.0 | 99.7 | 94.0 | 99.7 | 99.5 | 96.8 | 96.8 | 96.8 | 95.8 | 96.8 | 96.8 | 96.8 |
| HT_004 | 94.0 | 99.7 | | 93.8 | 99.5 | 93.8 | 99.5 | 99.7 | 97.0 | 97.0 | 97.0 | 96.0 | 97.0 | 97.0 | 97.0 |
| HT_084 | 99.2 | 94.0 | 93.8 | | 93.8 | 99.0 | 93.8 | 93.5 | 96.3 | 96.3 | 96.3 | 96.3 | 96.3 | 96.3 | 96.3 |
| HT_126 | 94.0 | 99.7 | 99.5 | 93.8 | | 93.8 | 99.5 | 99.2 | 96.5 | 96.5 | 96.5 | 95.6 | 96.5 | 96.5 | 96.5 |
| HT_128 | 99.7 | 94.0 | 93.8 | 99.0 | 93.8 | | 93.8 | 94.0 | 96.8 | 96.8 | 96.8 | 95.8 | 96.8 | 96.8 | 96.8 |
| HT_144 | 94.0 | 99.7 | 99.5 | 93.8 | 99.5 | 93.8 | | 99.2 | 96.5 | 96.5 | 96.5 | 95.6 | 96.5 | 96.5 | 96.5 |
| Sample1 | 93.8 | 99.5 | 99.7 | 93.5 | 99.2 | 94.0 | 99.2 | | 97.3 | 97.3 | 97.3 | 96.3 | 97.3 | 97.3 | 97.3 |
| Sample2 | 96.5 | 96.8 | 97.0 | 96.3 | 96.5 | 96.8 | 96.5 | 97.3 | | 100.0 | 100.0 | 99.0 | 100.0 | 100.0 | 100.0 |
| Sample3 | 96.5 | 96.8 | 97.0 | 96.3 | 96.5 | 96.8 | 96.5 | 97.3 | 100.0 | | 100.0 | 99.0 | 100.0 | 100.0 | 100.0 |
| Sample4 | 96.5 | 96.8 | 97.0 | 96.3 | 96.5 | 96.8 | 96.5 | 97.3 | 100.0 | 100.0 | | 99.0 | 100.0 | 100.0 | 100.0 |
| Sample5 | 95.6 | 95.8 | 96.0 | 96.3 | 95.6 | 95.8 | 95.6 | 96.3 | 99.0 | 99.0 | 99.0 | | 99.0 | 99.0 | 99.0 |
| Sample6 | 96.5 | 96.8 | 97.0 | 96.3 | 96.5 | 96.8 | 96.5 | 97.3 | 100.0 | 100.0 | 100.0 | 99.0 | | 100.0 | 100.0 |
| Sample7 | 96.5 | 96.8 | 97.0 | 96.3 | 96.5 | 96.8 | 96.5 | 97.3 | 100.0 | 100.0 | 100.0 | 99.0 | 100.0 | | 100.0 |
| Sample8 | 96.5 | 96.8 | 97.0 | 96.3 | 96.5 | 96.8 | 96.5 | 97.3 | 100.0 | 100.0 | 100.0 | 99.0 | 100.0 | 100.0 | |

**Figure S7: Sequence alignment of ITS-1 ASVs generated by amplicon sequencing from fecal samples of Côte d'Ivoire patients and ITS-1 sequences extracted from whole genome sequence data from the eight adult worms obtained through the expulsion study**

Panel A: The figure shows the multiple sequence alignment of the 406 bp long ribosomal ITS-1 sequences generated from *Trichuris* population in Côte d'Ivoire from amplicon sequencing of fecal samples and ITS-1 sequences extracted from WGS of adult worms. The dissimilarities are highlighted.

Panel B: Distance matrix displaying the percentage identity among the ITS-1 sequences.

A

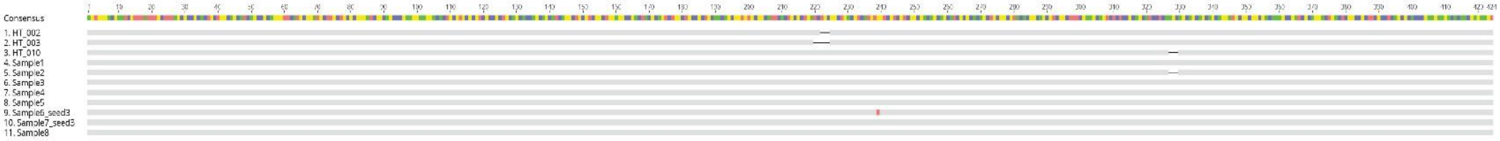


B

| | HT_002 | HT_003 | HT_010 | Sample1 | Sample2 | Sample3 | Sample4 | Sample5 | Sample6 | Sample7 | Sample8 |
|---|---|---|---|---|---|---|---|---|---|---|---|
| HT_002 | | 99.5 | 98.6 | 99.3 | 98.6 | 99.3 | 99.3 | 99.3 | 99.1 | 99.3 | 99.3 |
| HT_003 | 99.5 | | 98.1 | 98.8 | 98.1 | 98.8 | 98.8 | 98.8 | 98.6 | 98.8 | 98.8 |
| HT_010 | 98.6 | 98.1 | | 99.3 | 100.0 | 99.3 | 99.3 | 99.3 | 99.1 | 99.3 | 99.3 |
| Sample1 | 99.3 | 98.8 | 99.3 | | 99.3 | 100.0 | 100.0 | 100.0 | 99.8 | 100.0 | 100.0 |
| Sample2 | 98.6 | 98.1 | 100.0 | 99.3 | | 99.3 | 99.3 | 99.3 | 99.1 | 99.3 | 99.3 |
| Sample3 | 99.3 | 98.8 | 99.3 | 100.0 | 99.3 | | 100.0 | 100.0 | 99.8 | 100.0 | 100.0 |
| Sample4 | 99.3 | 98.8 | 99.3 | 100.0 | 99.3 | 100.0 | | 100.0 | 99.8 | 100.0 | 100.0 |
| Sample5 | 99.3 | 98.8 | 99.3 | 100.0 | 99.3 | 100.0 | 100.0 | | 99.8 | 100.0 | 100.0 |
| Sample6 | 99.1 | 98.6 | 99.1 | 99.8 | 99.1 | 99.8 | 99.8 | 99.8 | | 99.8 | 99.8 |
| Sample7 | 99.3 | 98.8 | 99.3 | 100.0 | 99.3 | 100.0 | 100.0 | 100.0 | 99.8 | | 100.0 |
| Sample8 | 99.3 | 98.8 | 99.3 | 100.0 | 99.3 | 100.0 | 100.0 | 100.0 | 99.8 | 100.0 | |

**Figure S8: Sequence alignment of ITS-2 ASVs generated by amplicon sequencing from fecal samples of Côte d'Ivoire patients and ITS-2 sequences extracted from whole genome sequence data from the eight adult worms obtained through the expulsion study**

Panel A: The figure shows the multiple sequence alignment of the 424 bp long ribosomal ITS-2 sequences generated from *Trichuris* population in Côte d'Ivoire from amplicon sequencing of fecal samples and ITS-2 sequence extracted from WGS of adult worms. The dissimilarities are highlighted.

Panel B: Distance matrix displaying the percentage identity among the ITS-2 sequences.

### Supplementary Table 1

**Mean Fecal Egg Count (FEC) of the samples**

The table displays the mean Eggs Per Gram (epg) of feces calculated from the quadruplicate Kato-Katz analysis for each sample from Côte d'Ivoire, Lao PDR, and Pemba.

| Population | Patient ID | Mean EPG |
|---|---|---|
| Côte d'Ivoire | BL-A4463 | 835 |
| Côte d'Ivoire | BL-A4572 | 201 |
| Côte d'Ivoire | BL-A4574 | 193.5 |
| Côte d'Ivoire | BL-A4617 | 255.5 |
| Côte d'Ivoire | BL-A4651 | 238 |
| Côte d'Ivoire | BL-A4654 | 1151.5 |
| Côte d'Ivoire | BL-B0133 | 132.5 |
| Côte d'Ivoire | BL-B0393 | 189.5 |
| Côte d'Ivoire | BL-B0704 | 230 |
| Côte d'Ivoire | BL-B1977 | 237 |
| Côte d'Ivoire | BL-B2032 | 152.5 |
| Côte d'Ivoire | BL-B6033 | 353.5 |
| Côte d'Ivoire | BL-B7235 | 163 |
| Côte d'Ivoire | BL-B7534 | 262 |
| Côte d'Ivoire | BL-B8460 | 384.5 |
| Côte d'Ivoire | BL-B8563 | 354.5 |
| Côte d'Ivoire | BL-B8744 | 110.5 |
| Côte d'Ivoire | BL-B8745 | 730.5 |
| Côte d'Ivoire | BL-B8746 | 383.5 |
| Côte d'Ivoire | BL-B8823 | 555.5 |
| Côte d'Ivoire | BL-B8835 | 381 |
| Côte d'Ivoire | BL-B9113 | 318.5 |
| Côte d'Ivoire | BL-B9674 | 243 |
| Lao PDR | BL-9625 | 699.5 |
| Lao PDR | BL-2445 | 506 |
| Lao PDR | BL-7181 | 253.5 |
| Lao PDR | BL-7180 | 261 |
| Lao PDR | BL-A1221 | 242.5 |
| Lao PDR | BL-A1244 | 224 |

| | | |
|---|---|---|
| Lao PDR | BL-7374 | 148 |
| Lao PDR | BL-A0421 | 186.5 |
| Lao PDR | BL-9754 | 123.5 |
| Lao PDR | BL-7208 | 192.5 |
| Lao PDR | BL-9624 | 139 |
| Lao PDR | BL-9753 | 128 |
| Lao PDR | BL-A1303 | 145.5 |
| Lao PDR | BL-7192 | 96.5 |
| Lao PDR | BL-A0988 | 130 |
| Lao PDR | BL-A0986 | 159 |
| Lao PDR | BL-1316 | 90.5 |
| Lao PDR | BL-A2774 | 95 |
| Lao PDR | BL-A0293 | 94 |
| Lao PDR | BL-9023 | 84.5 |
| Lao PDR | BL-A0944 | 84.5 |
| Lao PDR | BL-A0992 | 97.5 |
| Lao PDR | BL-7480 | 105 |
| Lao PDR | BL-A1581 | 73 |
| Lao PDR | BL-A1621 | 72.5 |
| Lao PDR | BL-A0884 | 72.5 |
| Lao PDR | BL-A3012 | 103.5 |
| Lao PDR | BL-A1391 | 97 |
| Lao PDR | BL-7528 | 68 |
| Lao PDR | BL-A1834 | 84.5 |
| Lao PDR | BL-A2775 | 75 |
| Lao PDR | BL-9622 | 91 |
| Lao PDR | BL-A0985 | 71.5 |
| Lao PDR | BL-A1261 | 100 |
| Lao PDR | BL-7065 | 97 |
| Lao PDR | BL-7415 | 54.5 |
| Lao PDR | BL-7008 | 88.5 |
| Pemba | BL-43 | 55.5 |
| Pemba | BL-52 | 133.5 |
| Pemba | BL-55 | 70 |
| Pemba | BL-56 | 170.5 |

| | | |
|---|---|---|
| Pemba | BL-58 | 79 |
| Pemba | BL-59 | 68.5 |
| Pemba | BL-90 | 85 |
| Pemba | BL-103 | 400 |
| Pemba | BL-104 | 80.5 |
| Pemba | BL-117 | 51 |
| Pemba | BL-144 | 72 |
| Pemba | BL-152 | 262.5 |
| Pemba | BL-314 | 214.5 |
| Pemba | BL-724 | 77.5 |
| Pemba | BL-1095 | 76 |
| Pemba | BL-1108 | 87.5 |
| Pemba | BL-1366 | 55.5 |
| Pemba | BL-1465 | 214 |
| Pemba | BL-1516 | 92 |
| Pemba | BL-2155 | 162.5 |
| Pemba | BL-2163 | 429 |
| Pemba | BL-2165 | 309.5 |
| Pemba | BL-2175 | 150 |
| Pemba | BL-2245 | 152 |
| Pemba | BL-3015 | 92 |
| Pemba | BL-3444 | 76 |
| Pemba | BL-4176 | 492.5 |
| Pemba | BL-713 | 802.5 |
| Pemba | BL-4336 | 243.5 |
| Pemba | BL-396 | 280 |
| Pemba | BL-3017 | 724.5 |

**<u>Supplementary Table S2</u>**

***Trichuris* sequences included from GenBank**

The *Trichuris* sequences that were used in the phylogenetic analysis of the ribosomal and mitochondrial loci are listed in the table.

| GenBank accession / Nomenclature | Host | Location | Marker | Reference |
|---|---|---|---|---|
| GQ301555 | Human | Cameroon | ITS1-5.8S-ITS-2 | *Ravasi DF et al., 2012* |
| KJ588159 | Human | Uganda | ITS-2 | *Ghai RR et al., 2014* |
| JN181817 | Human | Jamaica | ITS-2 | *Nissen S et al., 2012* |
| JN181820 | Human | Jamaica | ITS-2 | *Nissen S et al., 2012* |
| JN181850 | Human | Uganda | ITS-2 | *Nissen S et al., 2012* |
| AM992984 | Human | China | ITS-1-5.8S-ITS-2 | *Liu GH et al., 2014* |
| AM992997 | Human | China | ITS-1-5.8S-ITS-2 | *Liu GH et al., 2014* |
| KJ588071 | Human | Uganda | ITS-1 | *Ghai RR et al., 2014* |
| KP336484 | Vervet monkey | Italy | ITS1-5.8S-ITS-2 | *Cavallero S et al., 2015* |
| JF690945 | Lion-tailed macaque | Czech Rep. | ITS-2 | Unpublished |
| KJ588162 | Red colobus | Uganda | ITS-2 | *Ghai RR et al., 2014* |
| KT344830 | White-cheeked gibbon | China | ITS-1-5.8S-ITS-2 | *Xie Y et al., 2018* |
| MH390366 | White-cheeked gibbon | China | ITS-1-5.8S-ITS-2 | *Xie Y et al., 2018* |
| AM992999 | Pig | China | ITS-1-5.8S-ITS-2 | *Liu GH et al., 2014* |
| MG656442 | Pig | China | ITS-1 | Unpublished |
| MG656443 | Pig | China | ITS-2 | Unpublished |
| AJ249966 | Pig | Spain | ITS-2 | *Cutillas C et al., 2007* |
| AP017704 | Human | Japan | Mitochondrion | Unpublished |
| KT449826 | Human | Uganda | Mitochondrion | *Hawash M et al., 2015* |
| KT449824 | Hamadryas baboon | Denmark | Mitochondrion | *Hawash M et al., 2015* |
| KT449825 | Olive baboon | USA | Mitochondrion | *Hawash M et al., 2015* |
| KT449823 | Pig | Denmark | Mitochondrion | *Hawash M et al., 2015* |
| GU070737 | Pig | China | Mitochondrion | *Liu GH et al., 2012* |

| | | | | |
|---|---|---|---|---|
| KY368772 | Human | Denmark | Mitochondrion | *Søe MJ et al., 2018* |
| MN_ESP_MAL_CG_001 | Colobus Monkey | Spain | Mitochondrion | *Doyle S et al., 2022* |
| MN_ESP_MAL_CG_002 | Colobus Monkey | Spain | Mitochondrion | *Doyle S et al., 2022* |
| MN_CHN_GUA_LM_008 | Francois leaf monkey | China | Mitochondrion | *Doyle S et al., 2022* |
| MN_UGA_KAB_HS_005 | Human | Uganda | Mitochondrion | *Doyle S et al., 2022* |
| MN_DNK_COZ_PH_001 | Baboon | Denmark | Mitochondrion | *Doyle S et al., 2022* |
| MN_HND_SAL_HS_001 | Human | Honduras | Mitochondrion | *Doyle S et al., 2022* |
| MN_HND_OLA_HS_001 | Human | Honduras | Mitochondrion | *Doyle S et al., 2022* |
| MN_CHN_GUA_HS_001 | Human | China | Mitochondrion | *Doyle S et al., 2022* |
| MN_CHN_GUA_HS_005 | Human | China | Mitochondrion | *Doyle S et al., 2022* |
| NC_025751 | Coyote | USA | Mitochondrion | *Mohandas et al., 2014* |
| NC_025754 | Human | Thailand | Mitochondrion | *Mohandas et al., 2014* |
| NC_025749 | Raccoon | Russia | Mitochondrion | *Mohandas et al., 2014* |
| NC_002681 | N/A - Larvae | USA | Mitochondrion | *Lavrov and Brown, 2000* |
| MZ229684 | Vole | Czech Republic | Mitochondrion | *Petruzela et al., 2021* |
| JQ996231 | Yak | China | Mitochondrion | *Liu et al., 2012* |
| MZ229690 | Natal multimammate mouse | Tanzania | Mitochondrion | *Petruzela et al., 2021* |
| KY368775 | N/A - Latrine | Denmark | Mitochondrion | *Soe et al., 2018* |
| JQ996232 | Addax | China | Mitochondrion | *Liu et al., 2012* |
| MG189593 | Golden snub-nosed monkey | China | Mitochondrion | *Wang et al. 2019* |
| MK333462 | Sheep | China | Mitochondrion | *Ahmad et al., 2019* |
| MZ229685 | Greater bandicoot rat | Laos | Mitochondrion | *Petruzela et al., 2021* |

| | | | | |
|---|---|---|---|---|
| MZ229688 | Mahomet mouse | Ethiopia | Mitochondrion | *Petruzela et al., 2021* |
| MZ229686 | Guinea multimammate mouse | Ethiopia | Mitochondrion | *Petruzela et al., 2021* |
| KC461179 | Francois leaf monkey | China | Mitochondrion | *Liu et al., 2013* |
| MZ229689 | Misonne's soft-furred mouse | Kenya | Mitochondrion | *Petruzela et al., 2021* |
| MW448471 | Barbary macaque | Spain | Mitochondrion | *Rivero et al., 2021* |
| KT449822 | Pig | Denmark | Mitochondrion | *Hawash M et al., 2015* |
| GU385218 | Human | China | Mitochondrion | *Liu et al., 2012* |
| KY368765 | N/A - Latrine | Netherlands | Mitochondrion | *Soe et al., 2018* |
| KY368773 | N/A - Latrine | Denmark | Mitochondrion | *Soe et al., 2018* |
| MW448470 | Barbary macaque | Spain | Mitochondrion | *Rivero et al., 2021* |
| MW448472 | Guinea baboon | Spain | Mitochondrion | *Rivero et al., 2021* |

**Supplementary Table 3**

**Primers used for fecal stool DNA metabarcoding**

| | |
|---|---|
| Trichuris_ITS1_F | 5’-ACTGGCCGAACCAAGCCATC-3’ |
| Trichuris_ITS1_R | 5’-TCTACGAGCCAAGTGATCCAC-3’ |
| Trichuris_ITS2_F | 5’-ATGTCGACGCTACGCCTGTC-3’ |
| Trichuris_ITS2_R | 5’-TAGCCTCGTCTGATCTGAGG-3’ |
| Trichuris_COX1_F | 5’-TGGCATAGATATTGACACTCG-3’ |
| Trichuris_COX1_R | 5’-AAGCAGTCGTTGTAGTCTAAG-3’ |
| Trichuris_NAD1_F | 5’-ATACTTCAACCCGTCCTAGATG-3’ |
| Trichuris_NAD1_R | 5’-TCGAGTAAATCAAATGGAGTCC-3’ |
| Trichuris_NAD4_F | 5’-AGAGTCGGGATGTACTAGCGTA-3’ |
| Trichuris_NAD4_R | 5’-AGTCGGACCATTTACTTGAGTTC-3’ |

**Supplementary Table S4**
**Genetic diversity of the rDNA loci**

| Region | ITS-1 | | | | ITS-2 | | | |
|---|---|---|---|---|---|---|---|---|
| | N | $N_{ASV}$ | $H_d$ | π | N | $N_{ASV}$ | $H_d$ | π |
| Côte d'Ivoire | 21 | 7 | 0.52 | 0.001 | 15 | 3 | 0.56 | 0.000 |
| Laos | 34 | 92 | 0.98 | 0.003 | 26 | 32 | 0.82 | 0.006 |
| Pemba island | 29 | 63 | 0.96 | 0.003 | 23 | 13 | 0.69 | 0.006 |

N = Number of samples sequenced; $N_{ASV}$ = number of ASVs; $H_d$ = ASV heterozygosity; π = Nucleotide diversity

**Supplementary Table S5**
**Average pairwise nucleotide identity between *Trichuris* from Colobus monkeys (n=2), *T. suis* (n=3), and *T. hominis* (n=8) clades**

| | COX1 | | | All protein coding genes | | |
|---|---|---|---|---|---|---|
| | ***Trichuris sp.* Colobus Monkey** | ***T. suis*** | ***T. hominis*** | ***Trichuris sp.* Colobus Monkey** | ***T. suis*** | ***T. hominis*** |
| ***Trichuris sp.* ( Colobus Monkey)** | 99.6%* | 77.8% | 80.2% | 98.1%* | 71.6% | 73.1% |
| ***T. suis*** | 77.8% | 93.1%* | 78.9% | 71.6% | 91.7% | 71.7% |
| ***T. hominis*** | 80.2% | 78.9% | 99.4%* | 73.1% | 71.7% | 94%* |

*Indicates average pairwise nucleotide identity between samples of the same species / clade.
