## Appendix for "Discovery of a novel *Trichuris* species, *T. incognita*, with low sensitivity to albendazole-ivermectin treatment in Côte d’Ivoire patients, identified through fecal DNA metabarcoding" (from another version of this preprint: version 2, 2024-07-07)

### Appendix and Supplementary Tables

### Appendix A1: DNA extraction protocol for fecal samples

**Washing:**

1) Aliquot 500 µL (250 µL x2) of homogenized stool sample into the PowerBead Pro tube (from PowerFecal Pro kit). You should first remove the beads from the bead tube and transfer them to another tube before this step. (Aliquoting the sample in 2 x 250 µL volume ensures adequate homogenization and capturing as much stool as possible in the total 500 µL of starting material).
2) To the sample (500 µL) in the PowerBead Pro tube, add 1 mL molecular grade water, briefly vortex and spin in the mini centrifuge at 13,400 rpm (~12,000 *xg)* for five minutes using the Eppendorf Mini Spin Centrifuge..
3) Discard 500 µL of supernatant and add 500 µL ddH2O. Spin again at the same time and speed.
4) Discard 1000 µL of supernatant from the previous step. Do not add more water. Spin again for 2 minutes at the same speed.
5) After spinning, carefully discard around 250 - 300 µL of supernatant, depending on the nature and volume of your pellet.
6) After washing, you should now have a final volume of 200 µL - 250 µL of feces in water in your tube. Add 500 µL of CD1 solution (PowerFecal Pro kit), transfer the beads back to this tube, vortex well and carry out the optimization steps.

**The mechanical disruption steps are as follows:**

**x3**

1) Snap freeze the tubes by dipping them in liquid nitrogen for about 10 seconds or until they're completely frozen.
2) Heat at 100°C at 750 rpm for 15min in a thermoblock.
3) Use the Mini Bead Beater (MBB 96) to carry out bead beating for 3 minutes. Vortex after this step.
4) Repeat steps 1 to 3 three times.

**DNA extraction using the Powerfecal Pro Kit:**

1) After optimization, add the remaining 300 µL of CD1 solution, bringing the total volume of CD1 solution to 800 µL (The sample after optimization becomes very frothy, be careful while opening the caps and adding the remaining volume of CD1).

2) Secure the PowerBead Pro Tube horizontally on a Vortex Adapter and vortex at maximum speed for 10 min.

3) Centrifuge the PowerBead Pro Tube at 15,000 *x g* for **2 minutes** (centrifugation time changed compared to actual protocol).

4) Transfer the supernatant to a clean 2 mL Microcentrifuge Tube (provided). Transfer around **800 – 850 µL of supernatant** (depending on the nature and volume of the pellet).

5) Add 200 µL of Solution CD2 and vortex for 5 s.

6) Centrifuge at 15,000 *x g* for 1 minute. Avoiding the pellet, transfer up to **800 µL of supernatant** (depending on the volume of your pellet) to a clean 2 mL Microcentrifuge Tube (provided).

7) Add 600 µL of Solution CD3 and vortex for 5 seconds.

8) Load 650 µL of the lysate onto an MB Spin Column and centrifuge at 15,000 *x g* for 1 minute.

9) Discard the flow-through and repeat step 8 to ensure that all of the lysate has passed through the MB Spin Column.

10) Carefully place the MB Spin Column into a clean 2 ml Collection Tube (provided). Avoid splashing any flow-through onto the MB Spin Column.

11) Add 500 µL of Solution EA to the MB Spin Column. Centrifuge at 15,000 *x g* for 1 min.

12) Discard the flow-through and place the MB Spin Column back into the same 2 ml Collection Tube.

13) Add 500 µL of Solution C5 to the MB Spin Column. Centrifuge at 15,000 *x g* for 1 min.

14) Discard the flow-through and place the MB Spin Column into a new 2 mL Collection Tube (provided).

15) Centrifuge at up to 15,000 *x g* for 2 min. Carefully place the MB Spin Column into a new 1.5 mL Elution Tube (provided).

16) Add 100 µL of Solution C6 to the center of the white filter membrane. **Incubate at room temperature for 1 minute.**

17) Centrifuge at 15,000 *x g* for 1 min. Discard the MB Spin Column. Store the DNA in -20°C
